# Human cytomegalovirus gH/gL/gO binding to PDGFRα provides a regulatory signal activating the fusion protein gB that can be blocked by neutralizing antibodies

**DOI:** 10.1101/2025.01.08.631902

**Authors:** Eric P. Schultz, Lars Ponsness, Jean-Marc Lanchy, Matthias Zehner, Florian Klein, Brent J. Ryckman

## Abstract

Herpesviruses require membrane fusion for entry and spread, a process facilitated by the fusion glycoprotein B (gB) and the regulatory factor gH/gL. The human cytomegalovirus (HCMV) gH/gL can be modified by the accessory protein gO, or the set of proteins UL128, UL130 and UL131. While the binding of the gH/gL/gO and gH/gL/UL128-131 complexes to cellular receptors including PDFGRα and NRP2 has been well-characterized structurally, the specific role of receptor engagements by the gH/gL/gO and gH/gL/UL128-131 in regulation of fusion has remained unclear. We describe a cell-cell fusion assay that can quantitatively measure fusion on a timescale of minutes and demonstrate that binding of gH/gL/gO to PDGFRα dramatically enhances gB-mediated cell-cell fusion. In contrast, gH/gL/pUL128-131-regulated fusion is significantly slower and gH/gL alone cannot promote gB fusion activity within this timescale. The genetic diversity of gO influenced the observed cell-cell fusion rates, correlating with previously reported effects on HCMV infectivity. Mutations in gL that had no effect on formation of gH/gL/gO or binding to PDGFRα dramatically reduced the cell-cell fusion rate, suggesting that gL plays a critical role in linking the gH/gL/gO-PDGFRα receptor-binding to activation of gB. Several neutralizing human monoclonal antibodies were found to potently block gH/gL/gO-PDGFRα regulated cell-cell fusion, suggesting this mechanism as a therapeutic target.

**SIGNIFICANCE:** Development of vaccines and therapeutics targeting the fusion apparatus of HCMV has been limited by the lack an *in vitro* cell-cell fusion assay that faithfully models the receptor-dependent fusion characteristic of HCMV entry. The cell-cell fusion assay described here demonstrated that the binding of gH/gL/gO to its receptor, PDGFRα serves to regulate the activity of the fusion protein gB. Moreover, this regulatory mechanism is specifically vulnerable to inhibition by neutralizing antibodies. The cell-cell fusion assay described here provides a new tool to characterize neutralizing mAbs as therapeutic agents.

## INTRODUCTION

Human cytomegalovirus (HCMV) burdens the world through congenital infections causing cognitive delays and hearing loss, and reactivation of latent infections in immunocompromised transplant recipients and HIV/AIDS patients resulting in vascular diseases, graft rejection and other systemic diseases (1–4). At least 1 in 150 infants in the USA acquires HCMV infection *in utero* or shortly after birth via breastmilk and the associated health care costs during the first year of life are estimated at >$60K per infant (5). Congenital HCMV is overrepresented among non-whites of lower socioeconomic status, emphasizing HCMV as a health disparity (6). Accordingly, the development of safe and effective intervention approaches is a high priority. The live-attenuated (7–9) and adjuvanted-subunit (10, 11) vaccine candidates have all been based on single HCMV strains and have failed to exceed 50% efficacy. This seems to mirror the fact that naturally infected individuals can be “re-infected” by genetically distinct strains and this is associated with increased congenital infections in seropositive people (12–14).

Genomics studies have revealed deep complexities in the structure and dynamics of HCMV genetic diversity *in vivo* (15–23). Nineteen of the 165 canonical genes exist as multiple alleles, or “genotypes”. Due to high nucleotide (*nt*) diversity between alleles, these genes are often called “hyper-variable”, giving the impression of rapid, perpetual genetic drift as observed for RNA viruses. However, striking conservation within allele groups argues that the inter-allelic *nt* diversity is ancient and stable on a human timescale. Many of the prime vaccine targets are allelic, including the core glycoproteins involved in entry, gB, gH and gO. The remainder of the ∼235kb genome is comprised of conserved, mono-allelic genes that contain sporadic polymorphisms, and low linkage-disequilibrium, indicating frequent recombination that shuffles the allelic genes into a vast number of distinct haplotypes (16–18, 20, 21).

It is generally accepted that direct “cell-to-cell” spread is one way that viruses can evade the effects of neutralizing antibodies (nAb) (24). We have shown that in cell culture, genetically distinct strains of HCMV can have strong preferences for spread via diffusion of extracellular virus in the culture supernatant (i.e., “cell-free” spread), or by direct cell-to-cell spread (25). How HCMV spreads *in vivo* is less clear. Leukocyte depletion has been linked to reduced transmission of HCMV during blood transfusions, arguing against large amounts of infectious, cell-free HCMV in the blood (26, 27). While this is consistent with the model of hematogenous HCMV dissemination via monocyte/macrophages (28, 29), these were small scale studies that do not offer broad insights into roles of cell-free and cell-associated virus in other aspects of HCMV pathogenesis. The tendency of clinical isolates to display a cell-associated phenotype in culture does not necessarily indicate the global nature of HCMV *in vivo* since these observations can be influenced by the single cell-type monolayer cultures used and the specific strains isolated. Indeed, there are examples of clinical isolates that show cell-free phenotypes upon initial culturing (30, 31). It is also broadly appreciated that the major route of horizontal transmission is cell-free virus released in bodily fluids (32), suggesting that neutralizing mucosal IgA may offer protection. While cell-to-cell spread is generally considered less sensitive to inhibition by nAb than cell-free spread (24, 33), the mechanisms of cell-to-cell spread by HCMV are not sufficiently understood to conclude that nAb are irrelevant. Indeed, some nAbs seem to impede cell-to-cell spread *in vitro*, albeit less efficiently than for cell-free spread (25, 34). Finally, there is clinical evidence that nAbs against the gH/gL glycoprotein complexes of HCMV can offer protection against transplacental transmission and reactivation in transplant recipients (35–37). Neutralization is one likely mechanism driving this protection and is not mutually exclusive to others like Ab-dependent cellular cytotoxicity (ADCC) and Ab-dependent cellular phagocytosis (ADCP) (38, 39), and the entry mediating glycoproteins are key targets of nAbs.

Herpesvirus entry requires membrane fusion driven by glycoprotein gB under the regulation of gH/gL, and receptor-binding proteins like gD of herpes simplex virus (HSV) and gp42 of Epstein-Barr virus (EBV) (40, 41). The HCMV gH/gL can be bound by either gO, or the UL128-131 proteins, which act as receptor-binding domains. The gH/gL/gO complex binds to PDGFRα and this is required for efficient infection of fibroblasts (42, 43). Binding of gH/gL/pUL128-131 to receptors including NRP2 and OR14I1 facilitates infection of epithelial, endothelial and other select cell types (44, 45). There may be other receptors for gH/gL/gO since this complex is also important for infection of epithelial and endothelial cells, which may not express PDGFRα (43, 45–51). Three non-mututally exclusive mechanisms have been suggested for how these receptor-interactions facilitate infection: 1) virion attachment (47); 2) signal transduction influencing endocytic uptake or other cellular physiology (43, 44, 51, 52); and 3) regulation of the fusion protein, gB. While the latter mechanism is compelling by analogy with the action of gD for HSV and gp42 for EBV, no published data have yet directly linked receptor-binding by either gH/gL/gO or gH/gL/pUL128-131 to regulation of fusion.

Cell-cell fusion assays have been invaluable for studying herpesvirus entry (53–58). Transient expression of HCMV gB and gH/gL results in syncytia that develop slowly over 2-3 days (59, 60) reflecting the fundamental role of gH/gL as a regulatory co-factor for the fusion protein gB. However, the qualitative readout has precluded the use of syncytial cell-cell fusion assays to study the contribution of gO- or pUL128-131-receptor binding. Here we describe an improved HCMV cell-cell fusion assay based on split luciferase, similar that used by Anatasiu et al. to study HSV fusion (55). Our results confirm gH/gL as the core fusion co-factor for gB and demonstrate that binding of PDGFRα by gH/gL/gO provides receptor-dependent regulation of fusion, a mechanism that can be specifically targeted by nAbs.

## MATERIALS AND METHODS

### Cells lines

Retinal pigment epithelial cells (ARPE19) (American Type Culture Collection) were grown in a mixture of 1:1 DMEM and Ham’s F12 medium (DMEM-F12) (Sigma) supplemented with 10% FBS, penicillin-streptomycin, and amphotericin B. Primary human lung fibroblasts (MRC5; ATCC: CCL-171) were grown in DMEM supplemented with 6% heat-inactivated FBS and 6% BGS. 293IQ cells (Microbix, Toronto, Ontario, Canada) were grown in minimum essential medium (MEM; Life Technologies) supplemented with 10% FBS.

### Lentiviral and adenoviral vectors

The lckGFP or split GFP-RLuc_1-7_ gene was used to replace the enhanced green fluorescent protein (EGFP) open reading frame (ORF) in the pLJM1-EGFP lentiviral transfer vector plasmid. The pLJM1-EGFP plasmid was a gift from David Sabatini (Addgene plasmid no. 19319)(61). The plasmid was transformed in 293T cells together with three lentiviral helper plasmids. The pMDLg/pRRE, pRSV-Rev, and pMD2.G helper plasmids were a gift from Didier Trono (Addgene plasmid no. 12251, 12253, and 12259)(62). Two days after transformation, the lentiviral particles in the supernatant were purified from cell debris through syringe filtration and centrifugation. After titration, the particles were used to transduce low-passage ARPE19 cells. After a week of puromycin selection, cells were tested for RLuc1-7 expression by coinfection with split GFP-RLuc_8-11_ vectors, and aliquots were stored in liquid nitrogen until further use. Replication-defective (E1-negative) adenovirus (Ad) vectors that express HCMV TR gB, gH, gL, gO, PDGFRα-V242K, or Rluc_8-11_ were made as previously described(60). Briefly, Ad vector stocks were generated by infecting 293IQ cells at 0.1 PFU/cell for 6 to 10 days. The cells were pelleted by centrifugation, resuspended in DMEM containing 2% FBS, sonicated to release cell-associated virus, and cleared the cellular debris. Titers were determined by plaque assay on 293IQ cells. Multiplicities of infection (MOIs) for Ad vectors were determined empirically for each experiment and ranged from 3 to 30 PFU per cell.

### Fluorescence Microscopy

ARPE19 cells expressing membrane localized lckGFP were fixed with 4% paraformaldehyde and permeabilized using phosphate-buffered saline (PBS) containing 0.5% Triton X-100, 0.5% sodium deoxycholate, 1% bovine serum albumin (BSA), and 0.05% sodium azide. Cell nuclei were stained with 0.4 μM 4’,6‘-diamidino-2-phenylindole dihydrochloride (DAPI) as described previously (63).

### Syncytia Formation Assay

ARPE-19 expressing lckGFP cells were seeded in 12-well plates and allowed to grow to confluence, then the cells were infected with Ad vectors expressing the HCMV gB, gH, gL, gO, and PDGFRα-V242K proteins. Approx. 48-72 hours post infection, syncytia were analyzed by fluorescence microscopy.

### Real-time cell-cell fusion assay

ARPE19 cells constitutively expressing Rluc_1-7_ were plated in 96-well white-walled bioluminescence plates (Thermo) and transduced with adenovirus vectors encoding HCMV gB, gH, gL, and gO (or UL128, UL130, and UL131). Target cells (ARPE19 or MRC5) were transduced with Rluc_8-11_ and PDGFRα-V242K. 24 hours post transduction, effector cells were incubated with EnduRen live cell substrate for 1 hour at 37 deg C, then target cells were lifted with trypsin, resuspended in DMEM/F12 (no dye), and added to effector cells. Luminescence was measured every 10 min for 24 hours using a BioTek plate reader.

### CELISA

ARPE-19 epithelial cells were seeded in 96-well cell-based enzyme-linked immunosorbent assay (CELISA) culture plates (white wall, clear bottom) and transduced with Ad vectors expressing HCMV glycoproteins. To measure cell surface gH, cells were incubated for 1 h with 14-4b, fixed for 30 min, and then incubated for 45 min with secondary antibody. Cells were washed between steps with PBS supplemented with 1% BSA and 5% FBS. Two minutes prior to data collection, wells were incubated with SuperSignal enzyme-linked immunosorbent assay (ELISA) Femto substrate (Thermo), and then chemiluminescence was measured on a BioTek plate reader.

## RESULTS

### Binding of PDGFRα by gH/gL/gO provides positive regulation of the HCMV fusion protein, gB

To assess the role of PDGFRα-binding by gH/gL/gO in gB-mediated membrane fusion, retinal pigment epithelial cells (ARPE19) cells expressing plasma membrane-anchored GFP (lck-GFP) were transduced with adenovirus (Ad) expression vectors encoding HCMV glycoproteins and syncytia formation was assessed at 72 hours post transduction by fluorescence microscopy (Fig. 1). Consistent with the previous findings (59), gH/gL alone was sufficient to promote gB-mediated cell-cell fusion, regardless of coexpression with PDGFRα. Syncytia were also observed when cells expressed gH/gL/pUL128-131, independent of PDGFRα. However, no syncytia were observed when cells expressed gH/gL/gO unless PDGFRα was also expressed. This suggests that PDGFRα-binding is necessary for gH/gL/gO to promote gB-mediated cell-cell fusion.

**Figure 1.**
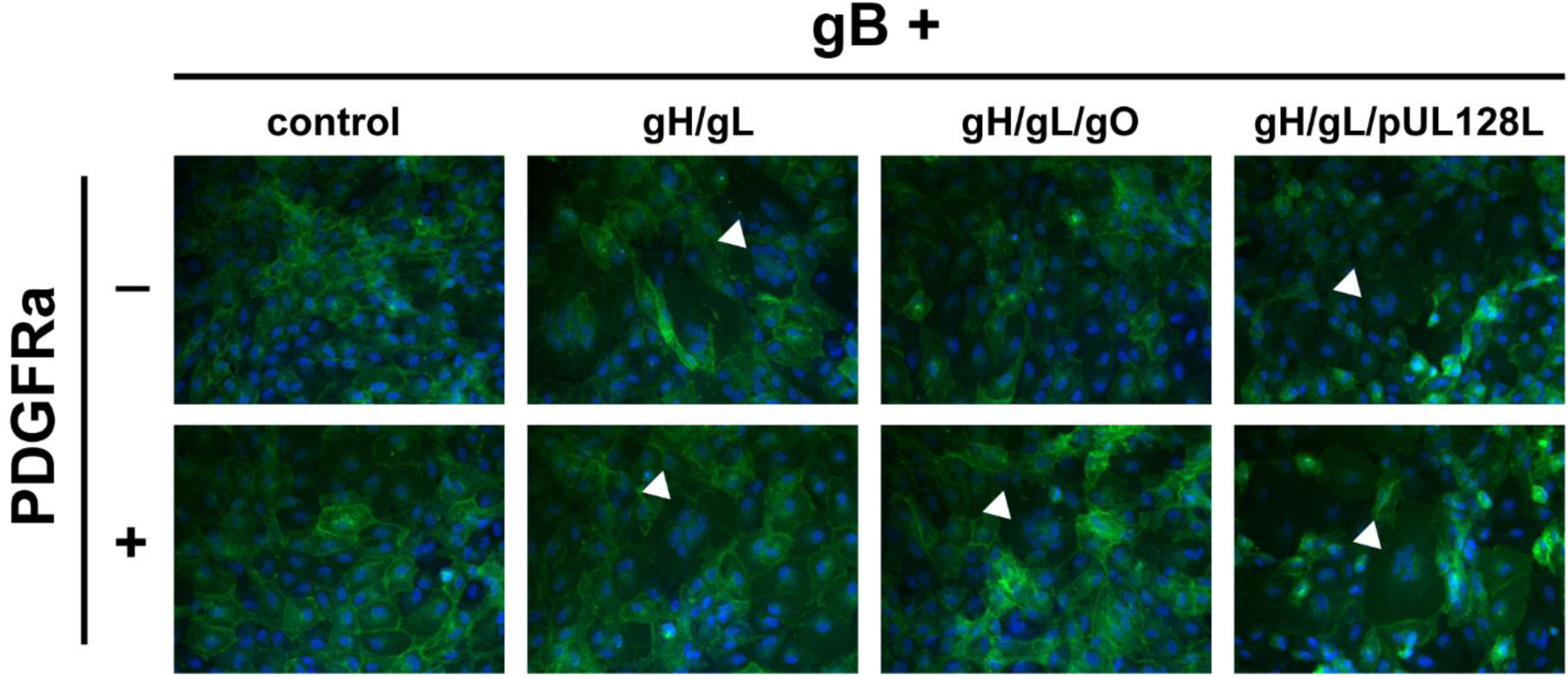
Syncytia formation resulting from co-expression of HCMV glycoproteins. ARPE19 cells stably expressing plasma membrane-anchored GFP(lck-GFP) were infected with adenovirus vectors encoding HCMV glycoproteins gB and gH/gL, gH/gL/gO, or gH/gL/pUL128-131 with or without PDGFRα. Nuclei were stained with DAPI and syncytia formation was monitored 48-72 h.p.i. by immunofluorescence. White arrows indicate representative syncytia for each condition.

To quantitatively compare the fusion resulting from the different combinations of HCMV glycoproteins, we used a live-cell, bimolecular complementation assay. Briefly, effector cells expressing one half of a GFP-rLuc protein (rLuc1-7) were transduced with Ad vectors encoding HCMV glycoproteins, preloaded with a cell-permeable luciferase substrate, and mixed with target cells expressing the other half (rLuc8-11). Cell-cell fusion was assessed by luminescence, recorded every 10 minutes for 20 hours (Figs 2A, B). Cell-cell fusion rates were determined by linear regression over the linear phase of each luciferase activity trace (Fig. 2C). When ARPE19 cells were used as effectors and targets, we observed more fusion with cells expressing gH/gL/pUL128-131 than those expressing gH/gL/gO or gH/gL alone (Fig. 2A). However, when PDGFRα-expressing ARPE19 cells were used as targets, dramatically more fusion was observed with gH/gL/gO-expressing effector cells (Fig 2B). The rate of fusion for gH/gL/gO effector cells was >100-fold higher with targets expressing PDGFRα over those without, and approx. 22-fold better than cells expressing gH/gL/pUL128-131, for which PDGFRα expression had no impact. The fusion rate for effector cells expressing gH/gL alone was indistinguishable from (-) gB control cells. However, the ability for gH/gL to promote syncytia formation over 2-3 days (Fig. 1) suggests an extremely low rate of fusion outside the timeframe of our quantitative assay.

**Figure 2.**
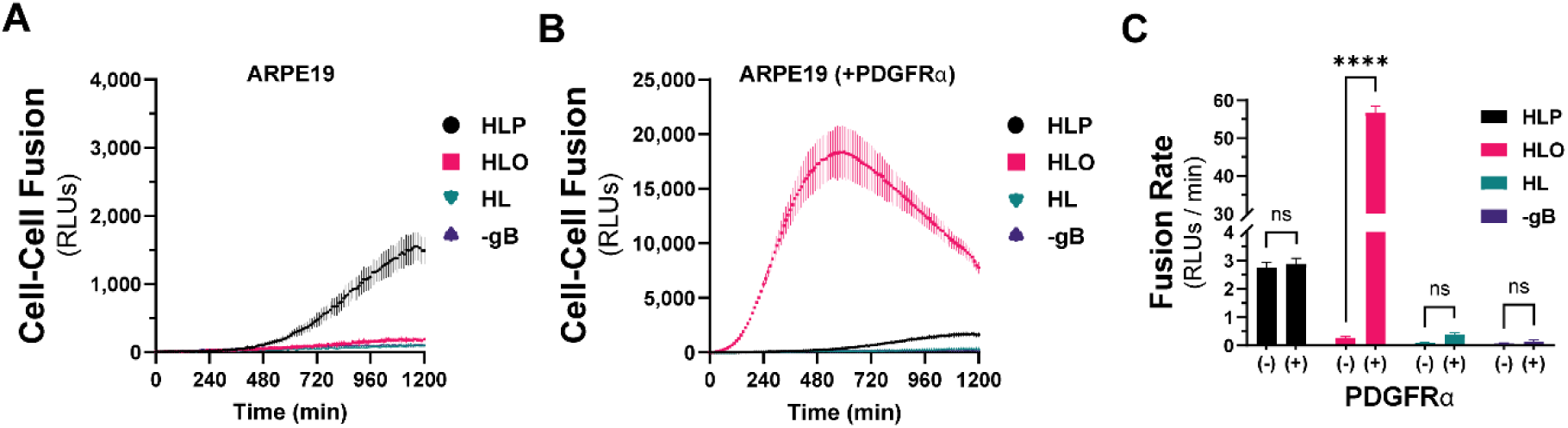
Quantitative assessment of HCMV fusion glycoproteins in real-time. A live-cell, biomolecular complementation assay (adapted from {}) for which target cells were added to effector cells expressing gB and gH/gL (HL), gH/gL/gO (HLO), or gH/gL/pUL128-131 (HLP) and luminescence was measured every 10 min for 20 hours. (A) Fusion traces for HCMV glycoprotein complexes with ARPE19 cells used as targets. (B) ARPE19 cells were infected with adenovirus encoding PDGFRα (V242K, {}) for 24 hours, then used as targets. (C) Fusion rates for all conditions were determined by linear regression over the linear phase of each luciferase activity trace. Error bars reflect the standard deviation of three experiments and p-values reflect 2-way ANOVA comparisons between PDGFRα +/- target cells (ns>0.05, * >0.01, ** >0.001, *** >0.0001, **** <0.0001).

To test whether the enhanced cell-cell fusion observed with gH/gL/gO and PDGFRα was due to increased surface levels of gH/gL/gO compared to gH/gL alone, we titrated gH/gL/gO surface expression over an 8-fold range, to levels comparable to gH/gL alone (Fig 3A). Cell-cell fusion rates were remarkably unaffected by reduced gH/gL/gO levels, with statistical significance only being achieved between the highest and lowest conditions (Fig 3B-C). Even with surface levels comparable to gH/gL/gO, gH/gL alone failed to fuse over control, suggesting its deficiency was not due to low surface expression. The insensitivity of fusion rate to the surface expression of gH/gL/gO was consistent with the insensitivity of HSV cell-cell fusion to the surface expression of gH/gL (55).

**Figure 3.**
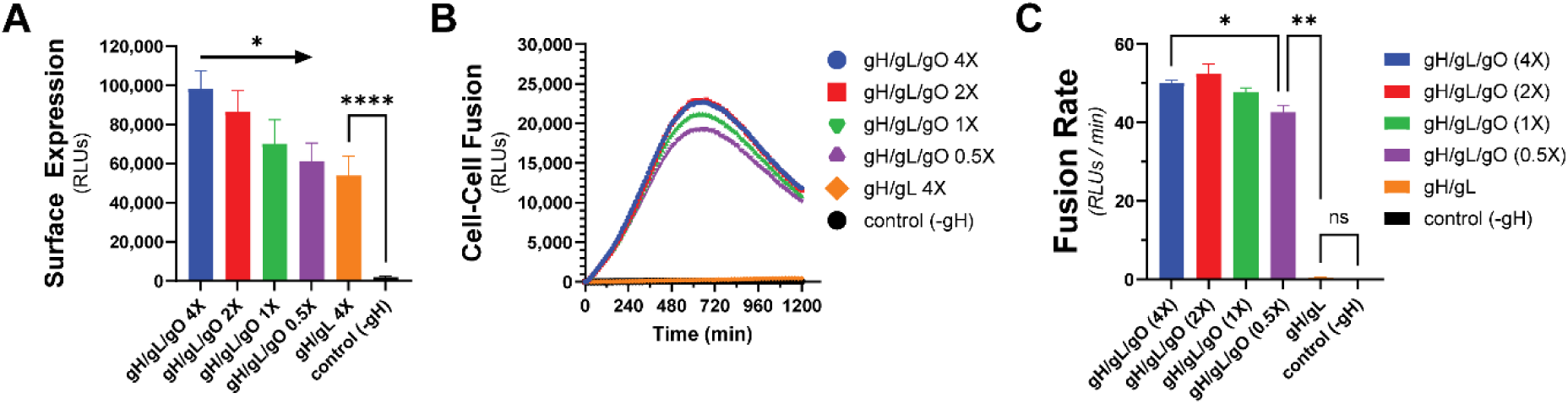
Fusion sensitivity to gH/gL/gO surface levels. The expression of gH/gL/gO was titrated by adjusting the adenovirus input over an 8-fold range, with 1X being the conditions used in Fig 2. (A) Surface expression was determined by CELISA using an antibody specific to gH (mAb 14-4b). (B) Fusion traces corresponding to the titrated gH/gL/gO levels. (C) Fusion rates corresponding to the titrated gH/gL/gO levels. Error bars reflect the standard deviation of three experiments and p-values were generated using ANOVA (ns>0.05, * >0.01, ** >0.001, *** >0.0001, **** <0.0001).

PDGFRα is expressed endogenously in most fibroblasts, with its highest expression in mesenchymal tissues including the lung, heart, intestine, skin and cranial facial mesenchyme (64). The lack of expression of PDGFRα in ARPE19 cells makes for an ideal cell-cell fusion system since it allows for a (-) PDGFRα control. To test whether endogenous levels of PDGFRα were sufficient for gH/gL/gO-regulated cell-cell fusion, we changed the target cell type in our assay to MRC5 fibroblasts. As was observed with PDGFRα-ARPE19 target cells, there was dramatically more fusion when the effector cells expressed gH/gL/gO compared to gH/gL/pUL128-131 or gH/gL alone (Fig. 4A, pink). Overexpression of PDGFRα in MRC5s led to significantly better fusion (Fig. 4A, brown), suggesting that cell-cell fusion is sensitive to the level of receptor on the surface of the target cell. Fusion of gH/gL/pUL128-131-expressing effectors cells with MRC5 targets cells was comparable to fusion with ARPE19 targets cells (compare Fig. 4B with 2C). In sum, these data support a model in which engagement of PDGFRα by gH/gL/gO provides an activation signal to regulate the fusion activity of gB.

**Figure 4.**
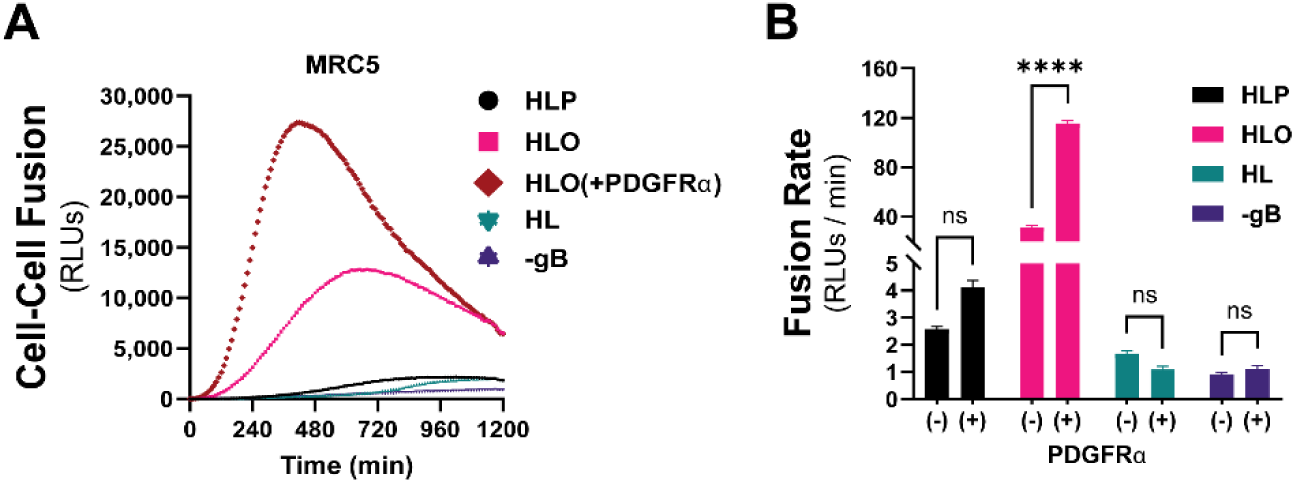
Regulation of gH/gL/gO-dependent fusion by endogenously expressed PDGFRα. (A) Fusion traces for HCMV glycoprotein complexes with MRC-5 cells used as targets. Fusion-regulation by gH/gL/gO was tested with endogenous levels (pink) and overexpressed (brown) PDGFRα. (B) Fusion rates for all HCMV glycoprotein complexes with endogenous (-) or overexpressed (+) of PDGFRα. Error bars reflect the standard deviation of three experiments and p-values were generated using ANOVA (ns>0.05, * >0.01, ** >0.001, *** >0.0001, **** <0.0001).

### Genetic diversity of gO can influence the kinetics of gH/gL/gO-regulated cell-cell fusion

The gene encoding gO, UL74, is one of several within the HCMV genome that show high levels of nucleotide diversity. Phylogenetic analyses indicate eight distinct alleles of UL74 gO with pairwise predicted amino acid differences among gO isoforms between 10-30% (65, 66). In Day et al., we reported a set of HCMV TR-based recombinants in which the endogenous gO allele (gO1b) was replaced with heterologous alleles (67). Among these, gO1a severely impaired virion infectivity, whereas gO2a gave a 30-fold enhanced infectivity and gO1c a modest 2-fold enhanced infectivity. To determine if these effects on infectivity were related to the role of gH/gL/gO in regulating gB, we compared these gO alleles in our quantitative cell-cell fusion assay (Fig 5A). While the differences in fusion rate were smaller than the corresponding differences in virus infectivity, the directionalities of the differences were congruent: as percent of parental gO1b: gO1a:83%; 1c:107%; 2a:112% (Fig. 5B). The differences in fusion rates could not be explained by differences in surface expression (Fig. 5C). This demonstrates that the diversity of gO can influence the kinetics of gH/gL/gO-PDGFRα-dependent fusion regulation, and this may contribute to observed infectivity differences among strains (46, 63).

**Figure 5.**
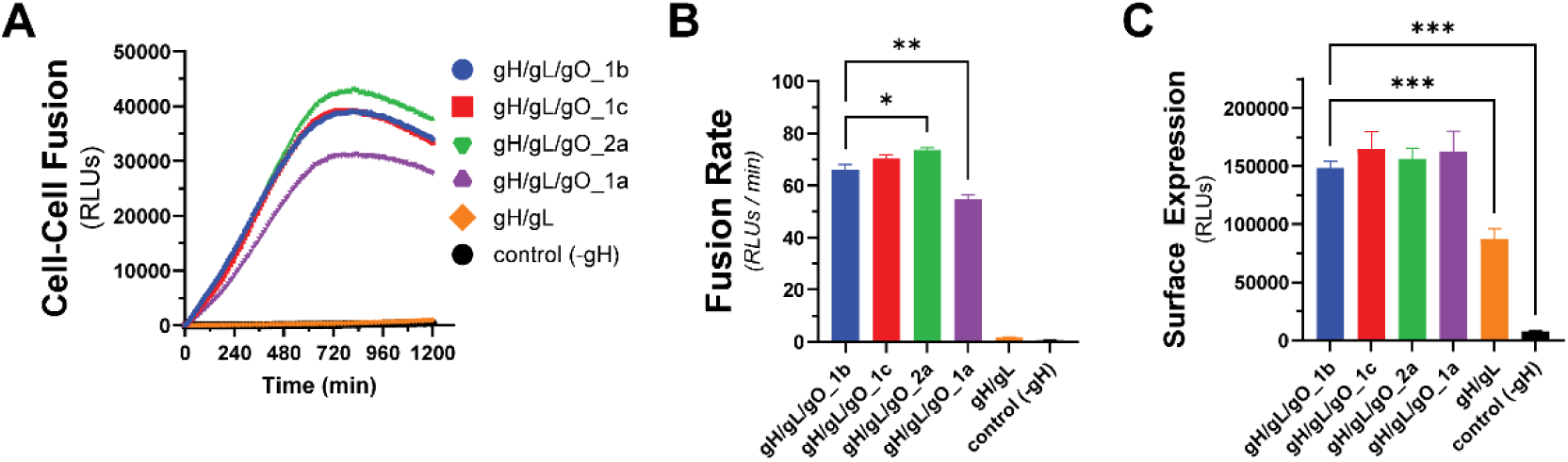
Effect of gO allele on gH/gL/gO-PDGFRα fusion regulation. (A) Real-time fusion traces for gH/gL/gO-dependent cell-cell fusion using 4 different gO alleles: 1b, 1c, 2a, 1a. (B) Fusion rates for different gO alleles. (C) Surface expression of gH/gL/gO with different gO alleles determined by CELISA. Error bars reflect the standard deviation of three experiments and p-values were generated using ANOVA (ns>0.05, * >0.01, ** >0.001, *** >0.0001).

### Kinetics of gH/gL/gO-PDGFRα dependent fusion regulation is sensitive to mutations in gL

We previously described a library of gL mutants that were able to form disulfide-linked gH/gL dimers and support assembly of gH/gL/pUL128-131 complexes capable of inducing receptor interference, but were unable to support the basal activity of gH/gL to promote gB-mediated cell-cell fusion (60). In a subsequent study, most of these gL mutants were shown to support stable soluble gH/gL/gO that could bind PDGFRα (68). Rescue of gL-null HCMV by most of these mutants resulted in moderately or severely reduced infectivity on fibroblasts, a gH/gL/gO-dependent parameter, while no effects were observed on gH/gL/pUL128-131-dependent aspects of HCMV infection. Here we analyzed a subset of these gL mutants for their ability to support the PDGFRα-dependent fusion regulation function of gH/gL/gO. Mutations L63, L139, and L201 reduced gH/gL/gO-dependent cell-cell fusion, roughly 15-fold, 2-fold, and 40-fold compared to WT gH/gL/gO, respectively, and L256 did not support fusion over WT gH/gL alone or even a condition lacking gH (Fig. 6). Given that the rate of fusion in this assay was insensitive to the surface expression of gH/gL/gO (Fig. 3), and none of these 4 mutations affected the binding of gH/gL/gO to PDGFRα (68), these data suggest gL is involved in the profusion signal post PDGFRα engagement to promote gB activation.

**Figure 6.**
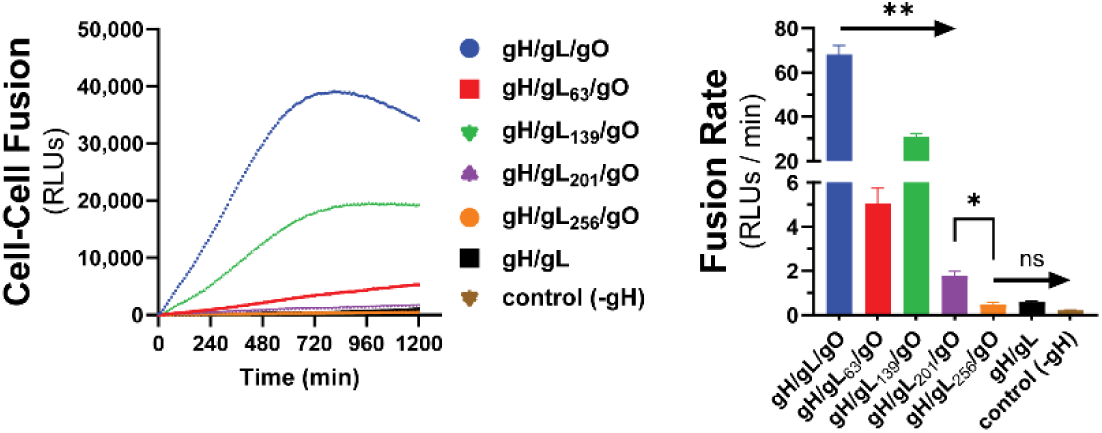
Effect of gL mutagenesis on gH/gL/gO-PDGFRα regulation of fusion. Real-time cell-cell fusion was performed using gL scanning alanine mutants. Fusion traces (A) and rates (B) are presented. Error bars reflect the standard deviation of three experiments and p-values were generated using ANOVA (ns>0.05, * >0.01, ** >0.001)

### Receptor-dependent regulation of fusion by gH/gL/gO is a target of antibody-neutralization

The gH/gL/gO complex is a major target of the humoral immune system with a plethora of antigenic domains including those that map to gH, defined by epitopes 13H11 and MSL109 (69), and others less well-defined on gO (70, 71). Zehner et al. isolated a set of 109 unique anti-gH/gL monoclonal Abs (mAbs) from the B-cell compartment of HCMV-infected donors that indicate at least 6 new antigenic groups distinct from those of 13H11 and MSL109 (72). These mAbs were characterized for their potency to neutralize two HCMV strains on fibroblasts, epithelial and endothelial cells, and for their ability to block binding of gH/gL/gO and gH/gL/pUL128-131 to PDGFRα or NRP2, respectively. Neutralization did not strictly correlate with blocking receptor-binding, indicating other neutralization mechanisms. To address the hypothesis that some of these Abs neutralize by blocking the receptor-dependent regulation of fusion by gH/gL/gO, a selection of nine Abs from this panel were tested for their ability to block gH/gL/gO-PDGFRα dependent cell-cell fusion (Fig. 7).

**Figure 7.**
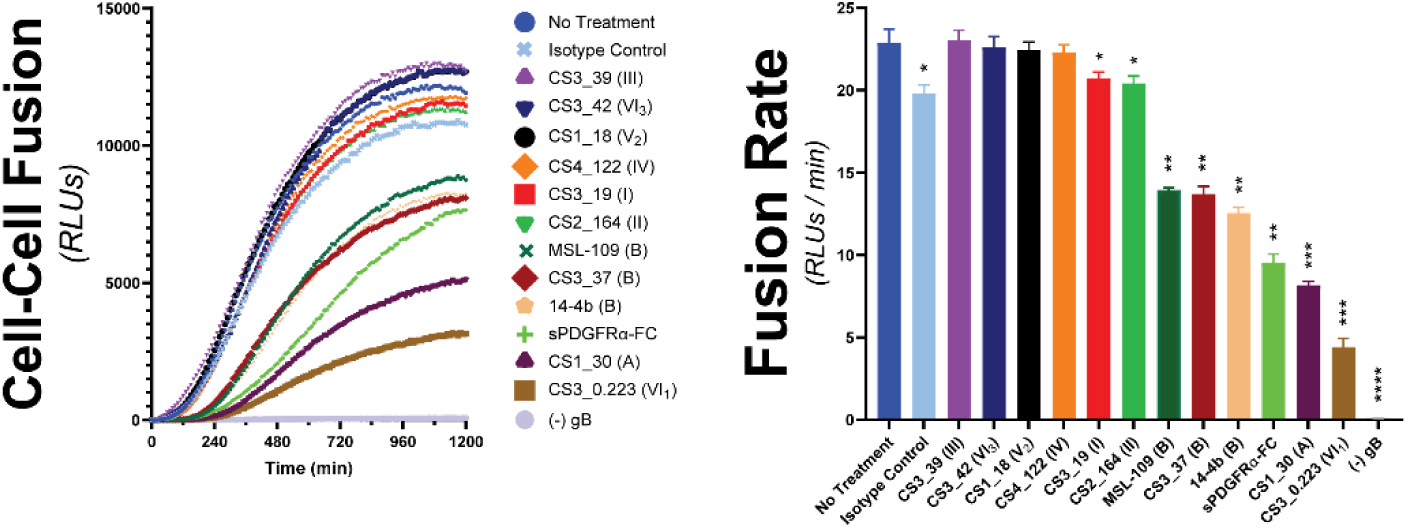
Assessing the ability for HCMV-neutralizing antibodies to block gH/gL/gO-PDGFRα regulation of fusion. The susceptibility of gH/gL/gO-dependent cell-cell fusion was assessed by preincubating effector cells with 50ug/mL of nAb for 1 hour prior to addition of target cells. Ab treatment was maintained and fusion was monitored over 20 hours. Several anti-gH/gL nAbs were tested including MSL-109(80), 14-4b(81), and eight novel Abs isolated from the B-cell compartment of HCMV-infected donors(72). Specific antigenic domains(72) are designated in parentheses. Fusion traces (A) and rates (B) are presented. Error bars reflect the standard deviation of three experiments and p-values were generated using ANOVA (ns>0.05, * >0.01, ** >0.001, *** >0.0001, **** <0.0001).

Six of the mAb clones tested failed to block cell-cell fusion over the isotype control (CS3_39, CS3_42, CS1_18, CS4_122, CS3_19, and CS3_164). The failure of CS3_39 and CS3_42 to block fusion was consistent with their failure to neutralize HCMV infection of fibroblasts, suggesting their ability to neutralize in epithelial and endothelial cells was due to blocking gH/gL/pUL128-131. In contrast, mAb clones CS1_18, CS4_122, CS3_19 and CS2_164 were neutralizing on fibroblasts but also failed to block cell-cell fusion, suggesting other neutralization mechanisms. Since soluble PDFGRα was able to inhibit cell-cell fusion, blocking gH/gL/gO binding to PDGFRα may be a plausible neutralization mechanism for CS3_19 and CS2_164. However, this may not always be sufficient for neutralization since CS3_39 neither neutralized virus, nor blocked cell-cell fusion despite being able to block PDGFRa-binding (72).

The mAbs tested that did block cell-cell fusion were each able to neutralize HCMV on fibroblasts, but did not block gH/gL/gO binding to PDGFRα (72). Ab CS3_37 inhibited cell-cell fusion rates comparably to MSL-109 and mAb 14-4b, approximately 2-fold. This was consistent with all three of these Abs belonging to the same antigenic group B (72), suggesting that the extent of inhibition may be linked to the specific region of gH/gL/gO targeted. The most potent inhibitors of cell-cell fusion were mAb clones CS1_30 and CS3_0.223, which reduced the fusion rate by 3-fold and 5.5-fold, respectively. For both, the inhibition was better than for soluble PDGFRα-FC, which could only reduce the fusion rate by 2.5-fold. mAb clone CS1_30 belongs to antigenic group A, defined by 13H11, but CS3_0.223 belongs to one of the novel antigenic groups and is currently unmapped (72). Together, these data support the notion that blocking gH/gL/gO-PDGFRα-dependent regulation of fusion may be a potent mAb neutralization mechanism.

## DISCUSSION

Cell-cell fusion assays have been used extensively as surrogates to study the fusion machinery of herpesviruses (53–58). For HCMV, transient expression of gB and gH/gL is sufficient to drive cell-cell fusion observed as syncytia (59). While this demonstrates the fundamental role of gH/gL as the direct cofactor for the fusion protein gB, it does not adequately model fusion during virus entry because 1) the syncytia formation takes 48-72 hours post transduction/transfection of gH/gL and gB expression constructs, and 2) *bona fide* HCMV entry requires either gO or pUL128-131 (48, 50, 73, 74). These accessory proteins serve as the receptor-binding subunits for gH/gL/gO and gH/gL/pUL128-131 (42, 44) but the specific role of these receptor interactions in facilitating infection have remained unclear.

Here, we adapted a live-cell, bimolecular complementation cell-cell fusion assay like that used by Anatasiu et al. to study the HSV fusion apparatus (55). This assay allows for precise discrimination of fusion kinetics over a wide dynamic range and on a timescale of minutes to hours, more representative of virus entry. Our results demonstrate that the binding of gH/gL/gO to its receptor PDGFRα provides a positive regulatory signal that activates the fusion protein gB. This mirrors the model for HSV fusion where binding of gD to any of several receptors, including Nectin-1 and HVEM, provides a signal or trigger to activate gH/gL as the cofactor for gB (reviewed in (41)). Expression of gH/gL/pUL128-131 also increased the fusion rate over that of gH/gL alone, but was dramatically lower than the rate observed with gH/gL/gO-PDGFRα, despite the target ARPE19 cells expression of the known pentamer receptors NRP2 and OR14I1 (44, 45). It is possible that the enhanced cell-cell fusion over gH/gL alone was secondary to increased cell surface expression resulting in more of the basal gH/gL cofactor activity (75, 76). Related to this, Vanarsdall et al. showed more syncytium formation in CD147-expressing HeLa cells when gH/gL/pUL128-131 was expressed compared to gH/gL alone (77). However, there was no evidence of direct interaction between gH/gL/pUL128-131 and CD147, so the mechanism of how CD147 promotes gH/gL/pUL128-131-dependent virus entry or cell-cell fusion was not clear. Thus, while our results demonstrate that binding of gH/gL/gO to PDGFRα serves a regulatory function for fusion, the specific function of receptor binding by gH/gL/pUL128-131 remains unclear. Finally, the fact that transient expression of PDGFRα was required for gH/gL/gO-dependent cell-cell fusion in ARPE19 cells indicates that these cells lack an endogenously expressed gH/gL/gO receptor on their surface, consistent with the endosomal route of entry into these cells (63).

The relationship of the measured cell-cell fusion rates to virus infectivity is not straightforward. Fusion rates measured using different alleles of gO corresponded to previously measured infectivity differences among heterologous gO allelic recombinant HCMV (Fig 5, (67)). However, the analysis of gL mutants revealed discrepancies. The gL mutations L139, L139, L201, L256 each reduced the cell-cell fusion rate and impaired the infectivity of HCMV TR, but L63 did not impact HCMV TR infectivity despite showing a reduced cell-cell fusion rate (Fig 6, (68)). Discrepancies in the magnitude of the measured effects may be partially explained by fundamental differences between the measurements including: 1) the cell-cell fusion assay measures a rate in real-time whereas infectivity is a static, endpoint parameter; 2) cell-cell fusion involves far more extensive membrane contacts than virus-cell fusion and may be less sensitive to receptor-binding differences; and 3) virus infection as measured by viral gene expression involves viral factors and cell process beyond those specifically related to fusion and the relative impact of fusion kinetics to virion infectivity may well be conditioned by these other factors.

The architectures of the L201 and L139 mutants seem consistent with the relative severity of their impacts on PDGFRα-dependent fusion regulation (69). The L201 mutation includes three arginine residues that lie in a groove between gH and gO (Fig. 8). R201 is 4.5Å from K252 of gO in the unbound gH/gL/gO structure, a distance that could be stabilized through interaction with a solvent ion, but these residues are rotated away from each other to 8.4Å when PDGFRα is bound. R207 is closely associated with a TYGRPI loop of gH in both the PDGFRα-bound, and unbound structures (Fig. 8, bottom right). The arginine of this gH loop also makes a salt-bridge contact with E51 of PDGFRα. R204 is not apparently involved in direct interactions, but the loss of this charged residue could influence the dynamics of the region and contribute to the loss of function. On the other hand, the L139 mutated residues make apparent interactions with gO but are quite distant from any gH or PDGFRα regions (Fig. 8, top right). Thus, the severe defect of the L201 mutation might reflect a critical role for this region in linking the binding of gH/gL/gO to PDGFRα to the regulation of gB. Further supporting this view, gO mutations near the L201 interface also impaired HCMV infectivity (48, 78).

**Figure 8.**
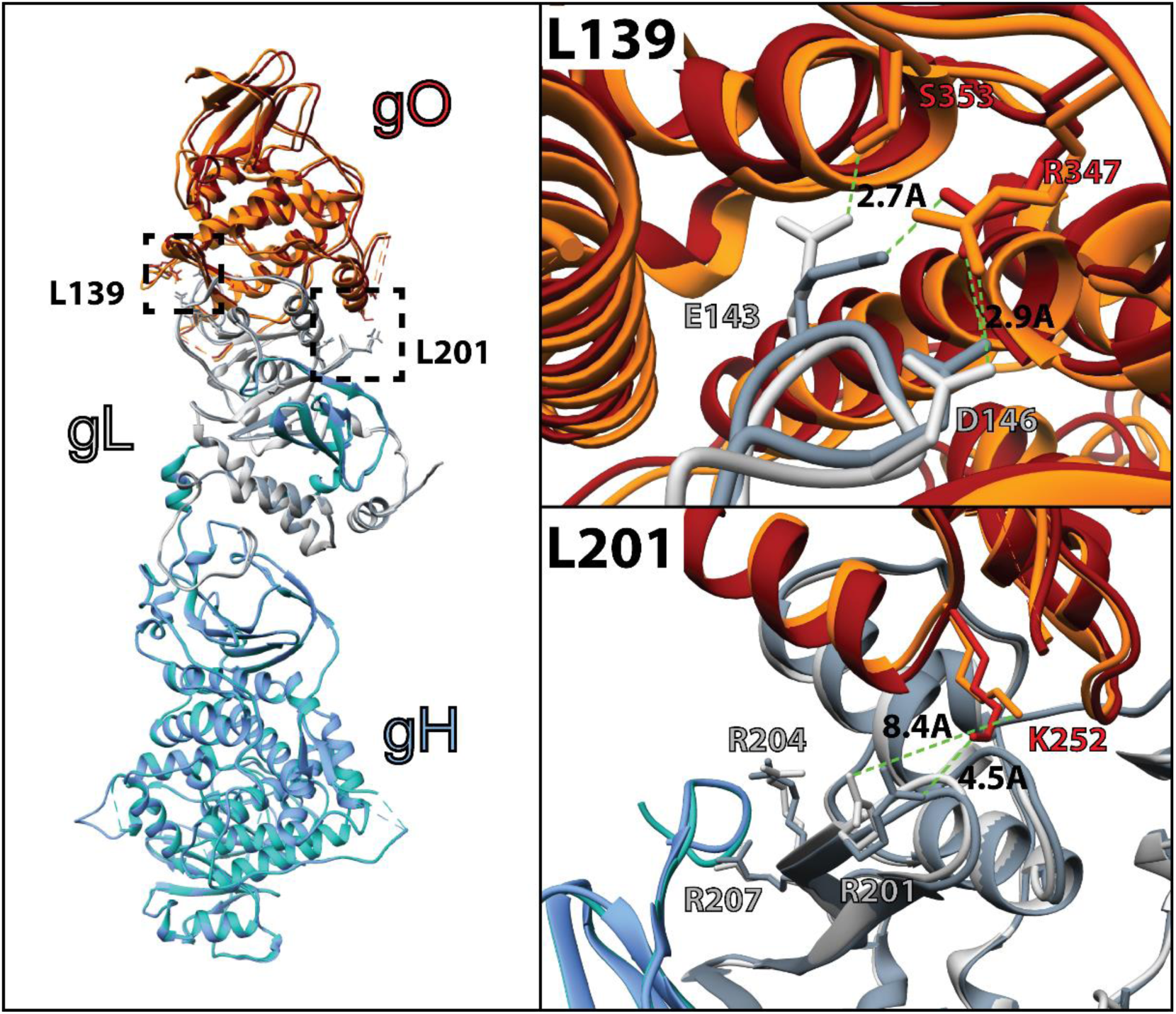
Comparison of the apo and PDGFRα-bound gH/gL/gO structure. CryoEM coordinates for the apo (pdb 7LBE) and PDGFRα-bound (7LBF) structures (69) of gH/gL/gO were aligned in Chimera (UCSF, (80)) and represented as ribbon structures. Individual subunits gH (blue), gL (grey), and gO (orange) are shaded to designate the apo (dark shade) and PDGFRα-bound (light shade). Residues of gL within the L139 (top right) and L201 (bottom right) regions that were mutated to alanine are displayed as sticks, with relevant interactions denoted with bond distances.

Receptor-dependent regulation of fusion represents a potential target mechanism for neutralizing Abs. The initial report of cell-cell fusion driven by gH/gL alone with gB showed that syncytium formation could be blocked with the neutralizing anti-gH mAb, 14-4b (59). Mutational analyses suggested that the 14-4b epitope overlaps with the defined MSL109 epitope at the membrane proximal region of gH and ELISA-based competition placed the CS3_37 epitope in the same antigenic group (60, 72). Consistent with this, all three of these Abs gave comparable inhibition of cell-cell fusion, suggesting a link between potency of inhibition and the specific antigenic domain. The most dramatic inhibition of cell-cell fusion was observed with the Ab CS3_0.223, which ELISA competition suggested reacts with novel antigenic domain, yet to be structurally defined (72).

Three of the nine novel anti-gH/gL mAb tested were shown to block the binding of gH/gL/gO to PDGFRα, but none of these inhibited cell-cell fusion (Table 1, (72)). This does not seem to indicate that inhibition of PDGFRα-binding fundamentally cannot inhibit cell-cell fusion since soluble PDGFRα was an effective inhibitor. Rather, this discrepancy may suggest that while soluble PDGFRα should be expected to exactly block the receptor-binding site on gH/gL/gO, receptor-blocking by anti-gH/gL Abs should be limited to steric hindrance imposed by Ab Fc domains and allosteric effects, which may be less effective. Moreover, it is possible that the interaction characteristics of soluble, immobilized gH/gL/gO and PDGFRα do not fully recapitulate those of the membrane bound versions of these proteins on the virus and cellular membranes such that these anti-gH/gL do not effectively block receptor-binding under physiologic conditions. Indeed, CS3_39 failed to neutralize HCMV at all, and neutralization by CS3_19 and CS2_164 may have been due to other mechanism such as virion aggregation or blocking adsorption (72). Thus, while inhibition of receptor binding is a plausible mechanism of neutralization, our results suggest that is not necessarily predictive of neutralization. The cell-cell fusion assay described here provides a new tool to characterize neutralizing mAbs.

**Table 1:** Inhibition characteristics of human anti-gH mAbs.

| mAb Clone<br>(antigen group) <sup>a</sup> | Cell-cell<br>fusion <sup>b</sup> | Receptor Binding <sup>a</sup> |  | HCMV Neutralization <sup>a</sup> |  |  |
| --- | --- | --- | --- | --- | --- | --- |
| | | PDFGRα | NRP2 | Fib | Epi | Endo |
| CS3_39 (III) | - | + | + | - | + | + |
| CS3_42(VI3) | - | - | - | - | + | + |
| CS1_18(V2) | - | - | - | + | + | + |
| CS4_122 (IV) | - | - | - | + | + | + |
| CS3_19 (I) | - | + | - | + | + | + |
| CS2_164 (II) | - | + | - | + | + | + |
| CS3_37(B) | + | - | - | + | + | + |
| CS1_30(A) | ++ | - | - | + | + | + |
| CS3_0.223(VI1) | +++ | - | - | + | + | + |

a. Zehner et al 2023,
b. Figure 7, herein

## ACKNOWLEDGMENTS

This work was supported by a grant from the National Institutes of Health (NIH) to B.J.R (R01AI097274), a fellowship from the American Heart Association (AHA) to E.P.S. (17POST33350043), a NIH CoBRE award to the Center for Biomolecular Structure and Dynamics at University of Montana (P30GM140963).

Experiments were designed by E.P.S., B.J.R., and J.-M.L. and performed by E.P.S., L.P., and the manuscript was prepared by B.J.R., E.P.S., M.Z and F.K.

